# Fertility is compromised after oocyte-specific deletion of Katanin A-subunit, *Katna1*, but not *Katnal1*

**DOI:** 10.1101/2023.10.22.563510

**Authors:** Wai Shan Yuen, Qing-Hua Zhang, Monique Dunstan, Deepak Aidhikari, Anne E O’Connor, Jessica EM Dunleavy, Moira K O’Bryan, John Carroll

## Abstract

Katanins are microtubule severing enzymes that play roles in shaping diverse microtubule-based structures during all cell cycle stages. To address the role of katanin A-subunits in mammalian oocytes, we have used the *Zp3*-*CreLox* approach to specifically delete katanin A1 (*Katna1*) and katanin A-like 1 (*Katnal1*) from the start of oocyte growth in mice. Here, we show that *Katnal1* is not required for normal female fertility, but that deletion of *Katna1* causes a 50% decrease in fertility. Further investigation in *Katna1*^-/-^ oocytes revealed no effect on MI spindle morphology but a significant effect on the morphology of MII spindles. This was accompanied by a decreased rate of fertilisation. Resultant *Katna1*^+/-^ heterozygous embryos that reached the 2-cell stage developed at normal rates to the blastocyst stage. Diploid homozygous parthenotes derived from *Katna1*^*-/-*^ oocytes revealed a reduced rate of blastocyst formation, decreased cell number and increased nuclear size. The ability of the paternal allele to rescue preimplantation development suggests the origin of the decrease in the fertility of conditional *Katna1*^*-/-*^ mice lies in abnormalities arising in the egg to embryo transition prior to embryonic genome activation.

## Introduction

The spindle is built in prometaphase of each cell cycle and is responsible for the accurate segregation of chromosomes in both meiotic and mitotic cell divisions. It is a highly dynamic structure with microtubules in a constant state of ‘flux’ driven by polymerisation at the chromosome-based kinetochore and depolymerisation at the spindle pole (Mitchison, 1989). Meiotic and mitotic cell cycles have evolved different mechanisms for building the spindle. In mitosis, spindle formation is typically centriole driven and relies on a ‘search and capture’ mechanism to position and align the chromosomes between the two centrioles that nucleate the spindle poles (Kirschner and Mitchison, 1986; Petry, 2016; Renda and Khodjakov, 2021). In female mammalian meiosis and early embryonic cleavage divisions, the absence of centrioles requires a chromosome mediated ‘inside-out’ approach to build the spindle (Blengini and Schindler, 2022; Brunet et al., 1998; Schuh and Ellenberg, 2007). The coordination of spindle formation and function relies on proteins that promote microtubule polymerisation and depolymerisation (Mihajlović and FitzHarris, 2018; Mogessie et al., 2018; Ohkura, 2015). An additional mode of microtubule regulation is microtubule severing, a process performed by the aptly named katanin family of proteins (McNally and Vale, 1993). Katanins have been implicated in spindle function in several cell types but their role in mammalian oocytes and embryos is largely unexplored.

Katanins are members of the ATPases Associated with diverse cellular Activities (AAA) protein superfamily and are highly conserved in eukaryotes, reflecting an essential role in contributing to microtubule organisation and function (McNally and Thomas, 1998; Srayko et al., 2000). Severing is ATP-dependent and occurs when the catalytic katanin A-subunit forms a hexameric ring that binds and severs microtubule filaments (Hartman, 1999; Hartman et al., 1998; Johjima et al., 2015; McNally and Vale, 1993). A regulatory subunit, katanin B-subunit typically enhances katanin-mediated severing, possibly via stabilisation of the katanin A-subunit hexamer (Dunleavy et al., 2021; Grode and Rogers, 2015), as well as playing a role in targeting it to the appropriate site, such as the centrosome or spindle pole (Hartman et al., 1998; McNally et al., 2000). In mammals, there are three katanin A-subunits, the canonical katanin p60 subunit A1 (KATNA1) in addition to paralogues KATNAL1 and KATNAL2, and two katanin B-subunits, the canonical katanin p80 subunit B1 (KATNB1) and paralog, KATNBL1 (McNally and Roll-Mecak, 2018). Microtubule severing is involved in both the remodelling and disassembly of existing microtubule structures and the generation of new microtubule structures, for example by generating short stable microtubule fragments for transport or seeding new growth, as well as by removing microtubules from nucleation sites (Ahmad et al., 1999; McNally et al., 2006; Nakamura et al., 2010; Srayko et al., 2006). Tubulin subunit extraction by microtubule severing proteins, such as katanin, has also been found to dynamically increase microtubule stability and amplification (Vemu et al., 2018). Consistent with these functions, KATNA1, KATNAL1, KATNAL2, and KATNB1 diffusely localise to cytoplasmic microtubules and concentrate at the centrosomes (Cheung et al., 2016; Hartman et al., 1998; McNally and Thomas, 1998; Ververis et al., 2016; Willsey et al., 2018).

Loss of function studies have revealed katanins are critical regulators of microtubule organisation in a wide range of cellular processes, including mitosis, cilia biogenesis, neurogenesis and spermatogenesis (Banks et al., 2018; Dunleavy et al., 2022; Dunleavy et al., 2021; Dunleavy et al., 2017; Hu et al., 2014; Lombino et al., 2019; Lynn et al., 2021; McNally and Roll-Mecak, 2018; Mishra-Gorur et al., 2014; O’Donnell et al., 2012; Smith et al., 2012). The loss of katanin activity in regulating the length and dynamics of microtubules leads to errors in cell division, shape, and migration (Buster et al., 2002; Lombino et al., 2019; O’Donnell et al., 2012; Smith et al., 2012). One of the best characterised roles for katanins is in male germ cell development, wherein *Katna1, Katnal1, Katnal2* and *Katnb1* are present in high levels, and knocking out these genes variously disrupts male meiosis, spermatid differentiation and Sertoli-germ cell adhesion (Dunleavy et al., 2022; Dunleavy et al., 2021; Dunleavy et al., 2017; O’Donnell et al., 2012; Smith et al., 2012).

In female meiosis, there is much less known. In *Caenorhabditis Elegans* oocytes, *Katna1* and *Katnb1* orthologues, MEI-1 and MEI-2, respectively are found to be essential for the normal assembly, function, cortical localisation and size of the meiotic spindle (Clark-Maguire and Mains, 1994; Mains et al., 1990; Srayko et al., 2000). In one particularly elegant study in *Xenopus* oocytes, katanin phosphorylation was found to control spindle size scaling (Loughlin et al., 2011). In *X. tropicalis, Katna1* lacks an inhibitory phosphorylation site at Serine 131 present in all other species including humans, leading to a particularly short spindle. By substituting *X. Tropicalis Katna1* with *X. Laevis Katna1* in egg extracts, spindle length was increased. The corollary was also true, spindle length was decreased in *X. Laevis* extracts by adding *X. Tropicalis Katna1*. Thus, katanins are critical in microtubule and spindle organisation in *C. Elegans* and *Xenopus* oocytes, but only limited evidence exists for a role in mammalian oocytes and embryos.

To this end, acute depletion of katanin A-like 1 (*Katnal1*), via in vitro transfection of siRNA in mouse oocytes, led to spindle disruption and inhibition of the two meiotic divisions (Gao et al., 2019). Here, we have developed an oocyte-specific Cre/LoxP model to investigate the *in vivo* function of KATNAL1, as well as the role of the highly conserved KATNA1 isoform in female meiosis. We find oocyte-specific deletion of *Katnal1* had no effect on female fertility with only modest effects on spindle structure. In contrast, oocyte-specific *Katna1* knockout females were found to be sub-fertile, with subsequent investigation revealing modest disruption of MII spindles and a decreased rate of fertilisation. Development of the resultant *Katna1*^+/-^ blastocysts *in vitro* was not affected, but a role for *Katna1* in cell division of preimplantation embryos was revealed in diploid *Katna1*^-/-^ parthenogenetic embryos lacking the paternal allele. Our findings demonstrate an evolutionarily conserved role of *Katna1* in female meiosis.

## Results

### *Katnal1*-deficient oocytes show normal spindle organisation and are fertile

Given *in vitro* studies indicate KATNAL1 regulates spindle function, we first turned to the ZP3-Cre-LoxP system to specifically deplete new KATNAL1 synthesis from the start of oocyte growth. This approach has been used extensively to generate fully grown oocytes lacking the protein of interest, in this case, KATNAL1. Reverse-transcription (RT) PCR across the deleted intron was used to confirm that no detectable *Katnal1* mRNA was present in fully grown oocytes, confirming the deletion was effective (Supplementary Fig. 1A-B).

To examine the role of KATNAL1 in MI spindle formation, oocytes were collected, and the MI oocytes were fixed at 8h post-IBMX release and their spindles examined for any abnormalities. Analysis of the *Katnal1*^*-/-*^ MI spindles showed significant, *albeit* marginal increase in spindle length (24.9μm versus 27.0μm; Fig 1D) and decrease in spindle width (15.7μm versus 15.0μm, Fig 1E). *Katnal1* deletion had no detectable effect on overall spindle morphology, as defined by barrel-shaped spindles with aligned chromosomes (96.3% versus 96.1%, Fig 1F). We next examined MII spindles in ovulated oocytes. In *Katnal*^*-/-*^ MII oocytes, spindles were sightly but significantly shorter (19.6μm versus 18.2μm; Fig 1G) and narrower (13.0μm versus 12.1μm, Fig 1H) compared to controls. Similar to MI oocytes, no significant difference was detected in overall spindle morphology (92.7% versus 87.7% normal, Fig 1I). Interestingly, 80% (11 out of 13) of abnormal spindles in the *Katnal1*^*-/-*^ MII oocytes present with microtubule asters throughout the cytoplasm (Supplementary Fig 1D), this was not seen in the abnormal control spindles.

**Figure 1.**
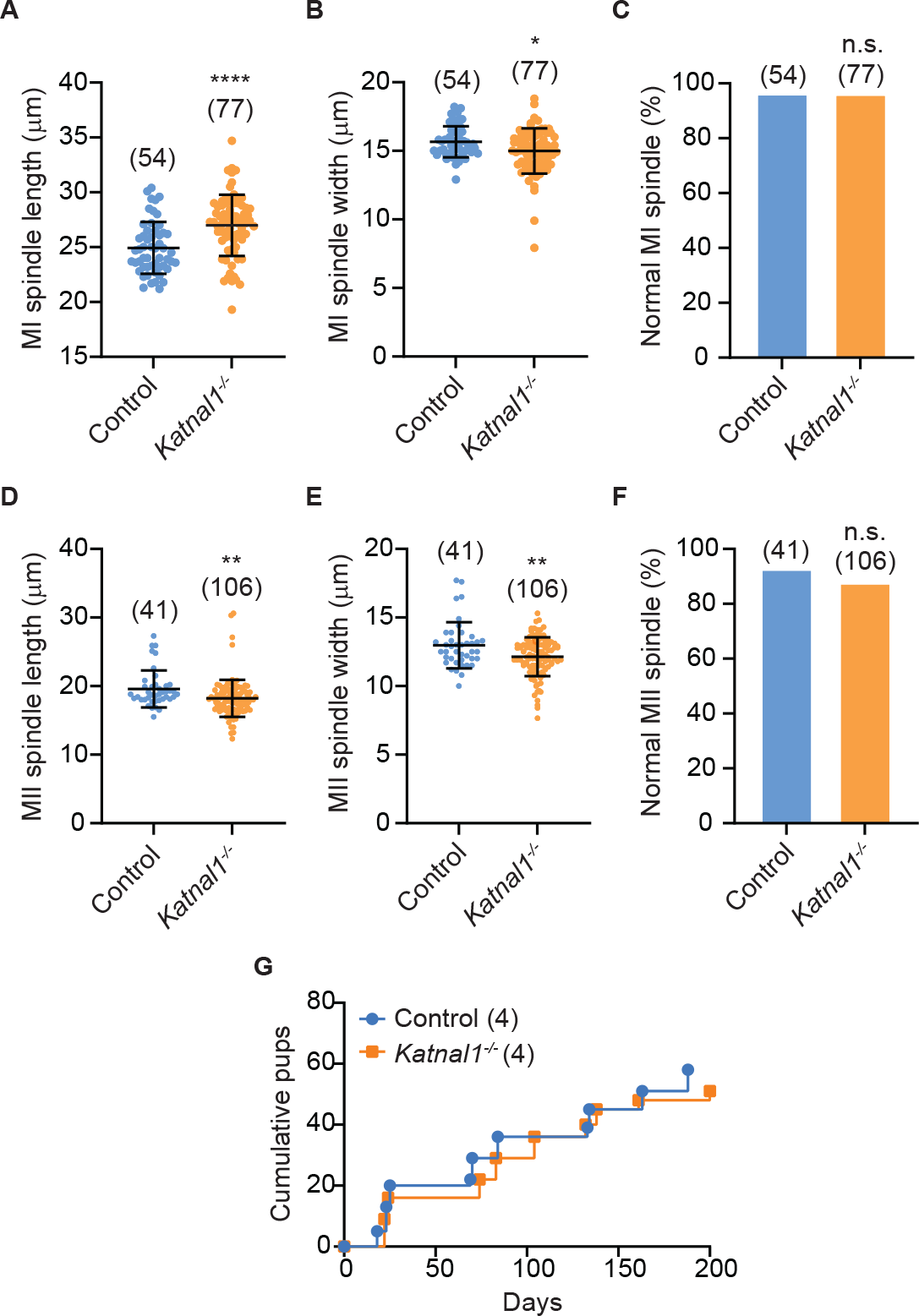
Katnal1 deficient mice present with minimal effects on fertility and female meiosis. (A) Spindle length of MI oocytes in control and Katnal1-/- groups. (B) Spindle width of MI oocytes in control and Katnal1-/- groups. (C) Percentage of MII eggs with normal spindle presentation in control and Katnal1-/- mice. (D) Spindle length of MII eggs in control and Katnal1-/- groups. (E) Spindle width of MII eggs in control and Katnal1-/- groups. (F) Percentage of MII eggs with normal spindle presentation in control and Katnal1-/- mice. (G) Cumulative pups born in control and Katnal1-/- mice over a 6-month period. Error bars indicate ± s.d. Scale bar, 10μm. Parentheses indict n numbers in each group. Statistical significance is depicted as n.s. p>0.05, ^**^p<0.0001.

Given the limited impact of *Katnal1* deletion on spindle organization, we tested for more subtle effects on fertility. However, when female conditional knock-out mice with *Katnal1*^*-/-*^ oocytes and *Katnal1*^*fl/fl*^ mice were mated with wildtype males over a period of 6 months we did not observe any effect on fertility (Fig 1G).

### *Katna1* deficient mice are subfertile and show evidence of MII spindle disruption

Given the lack of effect seen in oocyte-specific knockouts of *Katnal1*, we next examined the role of the canonical katanin A-subunit KATNA1 using the same Zp3-Cre-LoxP approach and confirmed the knockout in oocytes using RT-PCR (Supplementary Fig 1C). In contrast to *Katnal1*^*-/-*^ conditional knockouts, females with oocyte-specific deletion of *Katna1*^*-/-*^ showed a marked subfertility when mated with wildtype males for a period of 6 months (Fig 2A) – manifest as significantly smaller litter sizes in females with *Katna1*^*-/-*^ oocytes (Fig 2B, C).

**Figure 2.**
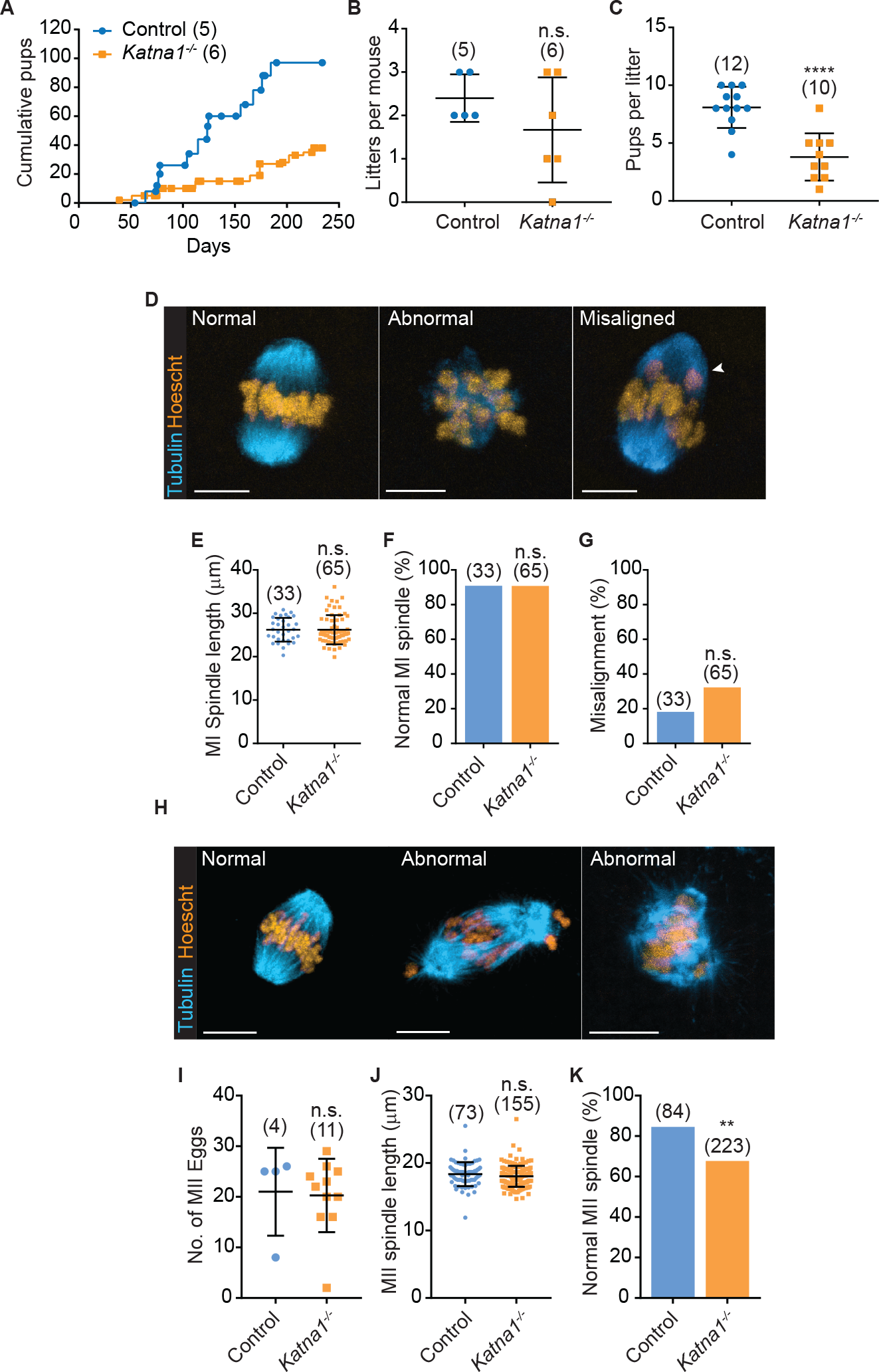
Mice with Katna1 deficient oocytes are subfertile. (A) Cumulative pups born in control and Katna1-/- mice over a 6-month period. (B) Total number of litters per mouse in control and Katna1-/- groups. (C) Number of pups in each litter birthed by control and Katna1-/- mice. (D) Representative images of normal, abnormal and misaligned MI spindle stained with -tubulin (blue) and Hoechst (orange). (E) MI spindle length of control and Katna1-/- mice. (F) Percentage of MI oocytes with normal spindles in control and Katna1-/- groups. (G) Chromosomal misalignment in MI oocytes of control and Katna1-/- mice. (H) Representative images of normal and abnormal MII spindle stained with -tubulin (blue) and Hoescht (orange). (I) No. of MII eggs collected from control and Katna1-/- mice. (J) Average MII spindle length of MII eggs retrieved from control and Katna1-/- groups, excluding those that present with non-bipolar spindles. (K) Percentage of MII eggs with normal spindle presentation in control and Katna1-/- mice. Scale bar, 10 m. Parentheses indict n numbers in each group. Statistical significance is depicted as n.s. p>0.05, ^*^p<0.0001.

To investigate whether the subfertility phenotype was caused by meiotic defects, we collected GV stage oocytes and examined spindles 8h after IBMX release (Fig 2D). MI stage *Katna1*^*-/-*^ oocytes showed no difference in spindle length (Fig 2E), overall morphology (Fig 2F), or alignment of chromosomes (Fig 2G). We went on to examine MII spindles in ovulated oocytes (Fig 2H) and no differences were seen in the number of oocytes ovulated in response to superovulation (Fig 2I), or in the length of MII spindles (Fig 2J). However, *Katna1*^*-/-*^ oocytes did show an increase in abnormalities in MII spindle morphology including misalignment of chromosomes and a disrupted barrel shape with unfocussed or absent spindle poles (15.5% versus 32.3%; Fig 2 H, K). This indicates KATNA1 is among a small number of proteins that appear to specifically regulate the MII spindle and play no role in MI.

### Fertilisation and development of *Katna1* deficient oocytes is compromised

The events of fertilisation and subsequent mitotic cell divisions may also be susceptible to the loss of KATNA1. Following *in vitro* fertilisation (IVF), a significant decrease was observed in proportions of *Katna1*^*-/-*^ oocytes that cleaved to the 2-cell stage compared to controls (72.1% versus 82.1%, p<0.05, Fig 3Ai). However, when these heterozygous *Katna1*^*+/-*^ 2-cell embryos were cultured to the blastocyst stage, the rate of blastocyst formation was not significantly different to controls (50.4% versus 54.0%; Fig 3Aii). Analysis of the heterozygous *Katna1*^*+/-*^ blastocysts using Imaris-based 3-D reconstruction was performed after staining for DNA and tubulin (Fig 3B). This analysis revealed a decreased blastocyst diameter (Fig 3C), although the average cell number (Fig 3D) and nuclear volume (Fig 3E) was unaffected. The length of spindles in mitotic cells of blastocysts derived from *Katna1*^*-/-*^ oocytes was increased (Fig 3F, G), but there was no difference in the proportion of mitotic cells in each blastocyst (Fig 3H).

**Figure 3.**
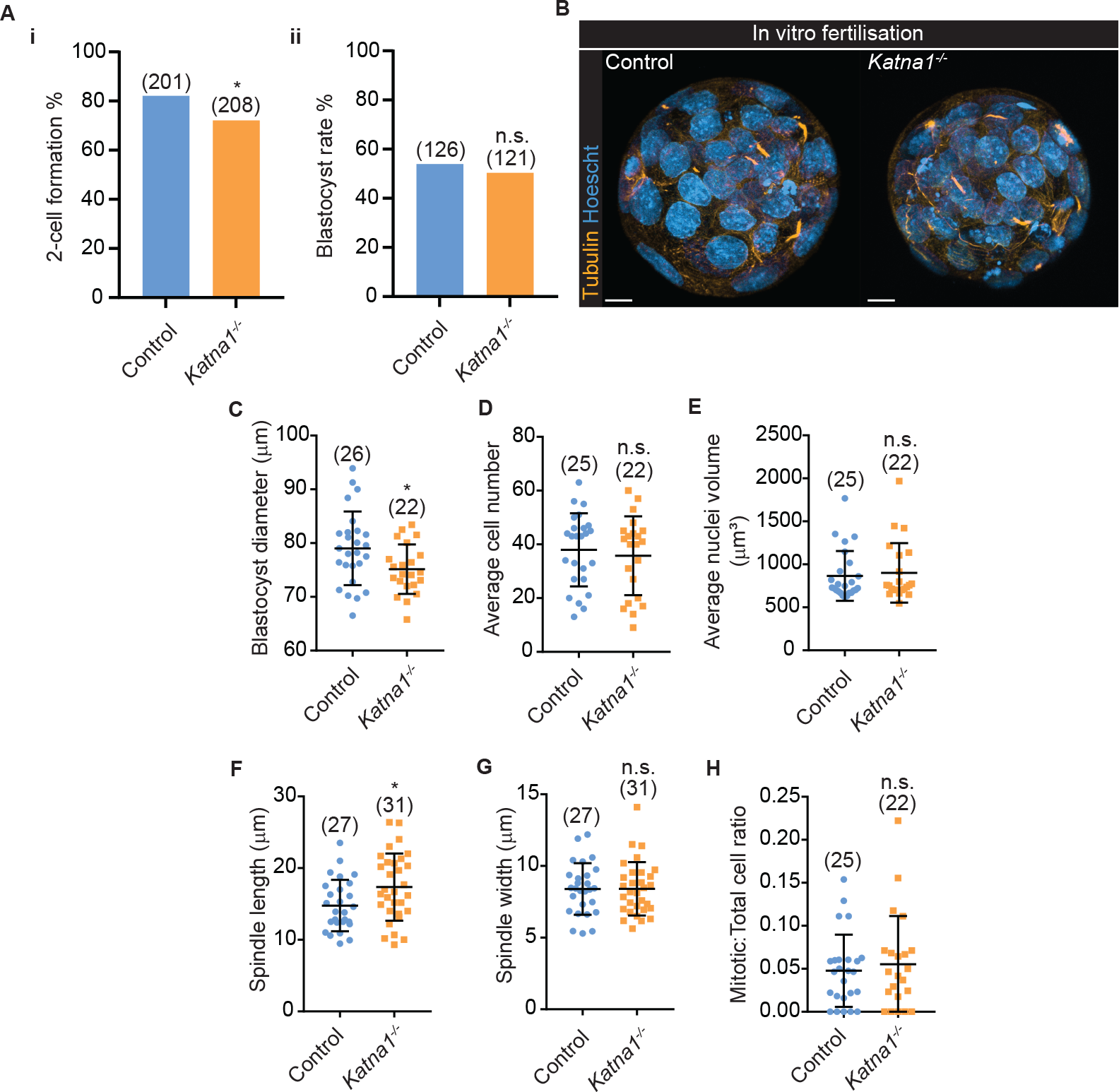
Fertilised Katna1 KO oocytes result in aberrant early embryogenesis. (Ai) Fertilisation rates of control and Katna1-/- mice from in vitro fertilisation. (Aii) Blastocyst formation rates of control and Katna1-/- groups. (B) Representative images of blastocysts collected at 4 days post IVF stained for α-tubulin (orange) and DNA (blue). Scale bar = 10μm. Aspects of control and knockout blasto-cysts from (B) were measured for; (C) average blastocyst diameter; (D) average cell number; (E) average nuclei volume; (F) spindle length; (G) spindle width; (H) mitotic cell to total cell number ratio. Parentheses indict n numbers in each group, pooled from at least two experimental replicates. Error bars indicate ± s.d. Statistical significance is depicted as n.s. p>0.05, ^*^p<0.05.

### Normal preimplantation development requires a functional *Katna1* allele

Fertilisation introduces a wildtype paternal *Katna1* allele which may serve to rescue any phenotypes beyond the 2-cell stage when the embryonic genome is activated. To better understand the role of KATNA1 in mitotic cell division in preimplantation embryos, we used parthenogenetic activation to create diploid homozygous *Katna1*^*-/-*^ embryos. As expected, similar proportions of *Katna1*^*-/-*^ and control oocytes formed 2-cell embryos after parthenogenetic activation (64.0% versus 75.0%; Fig 4Ai). However, the rate of blastocyst formation was dramatically reduced in *Katna1*^*-/-*^ embryos compared to controls (46.9% versus 93.9%; Fig 4Aii). Parthenogenetic blastocysts were analysed as described above for fertilised embryos (Fig 4B). The *Katna1*^*-/-*^ parthenogenetic blastocysts had a significantly reduced diameter (Fig 4C) lower mean cell number (Fig 4D), increased nuclear volume (Fig 4E), and increased spindle length, but no difference in spindle width (Fig 4G, H) compared to controls. Similar to IVF blastocysts, no difference was seen in the proportion of mitotic cells in each blastocyst (Fig 4F). Thus, KATNA1 is an important player in regulating microtubule function during preimplantation, but maternal effects of *Katna1* deletion can be rescued to some degree by the paternal allele introduced at fertilization.

**Figure 4.**
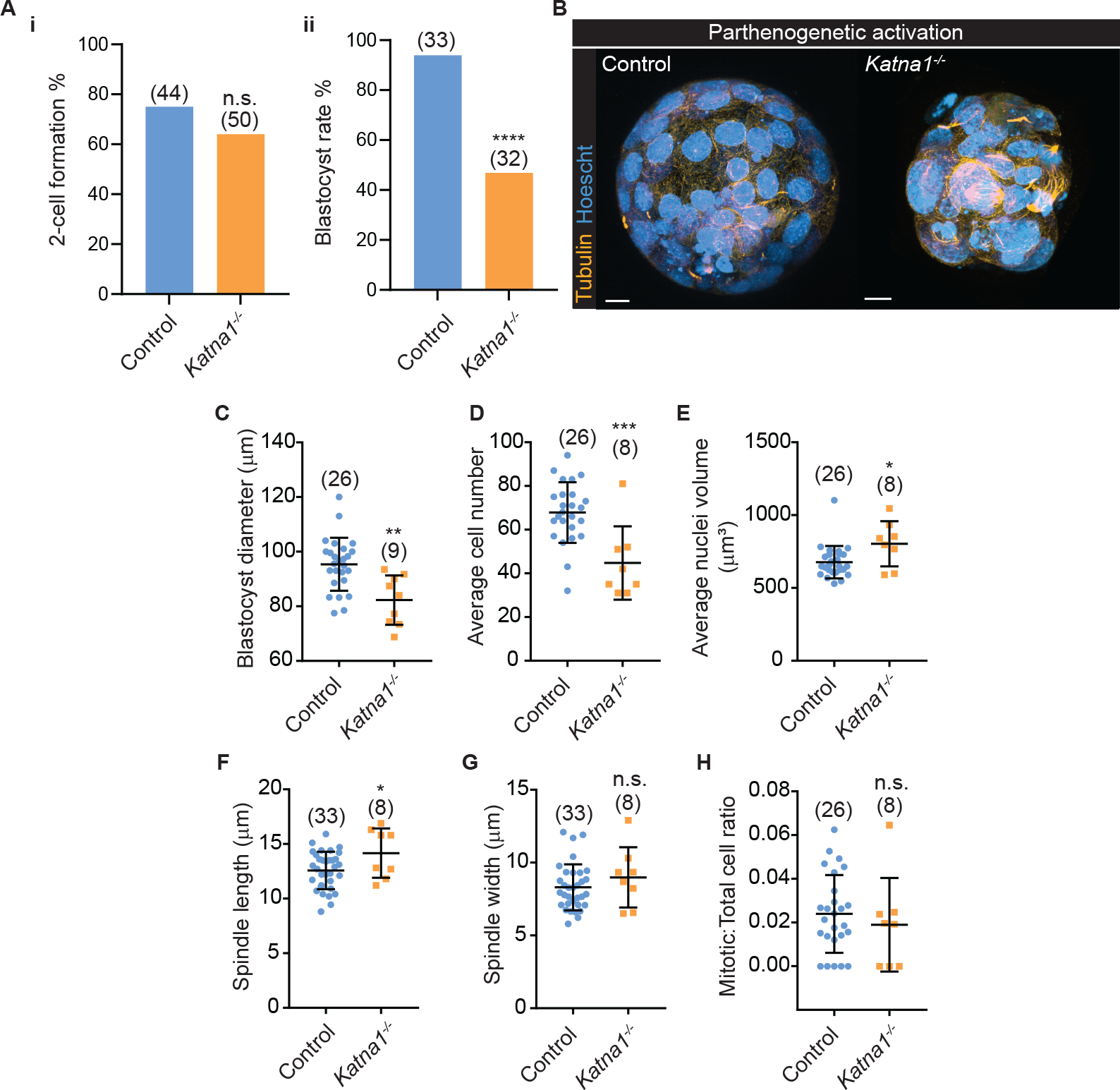
Katna1-deficient parthenotes exacerbate defects in blastocyst formation. (Ai) Fertilisation rates of control and Katna1-/- mice from parthenogenetic activation. (Aii) Blastocyst formation rates of control and Katna1-/- groups. (B) Representative images of blastocysts collected at 4 days post activation stained for α-tubulin (orange) and DNA (blue). Scale bar = 10μm. Aspects of control and knockout blastocysts from (B) were measured for; (C) average blastocyst diameter; (D) average cell number; (E) average nuclei volume; (F) spindle length; (G) spindle width; and (H) mitotic cell to total cell number ratio. Parentheses indict n numbers in each group, pooled from at least two experimental replicates. Error bars indicate ± s.d. Statistical significance is depicted as n.s. p>0.05, ^*^p<0.0001.

## Discussion

In lower order species *C. elegans* and *Xenopus*, katanin present in these species is well established as an essential regulator of female meiosis (Clark-Maguire and Mains, 1994; Loughlin et al., 2011; Mains et al., 1990; McNally and Thomas, 1998; Srayko et al., 2000). Herein, we reveal that in mammalian female meiosis, the canonical katanin A-subunit KATNA1 is a key microtubule regulator in MII but not MI oocyte spindles, in addition to having essential functions in the preimplantation embryonic mitoses that are required for embryo viability. Our data also shows that the non-canonical katanin A-subunit KATNAL1 assists in regulating spindle size and length in mouse oocytes but is ultimately dispensable for female fertility.

Oocyte-specific *Katna1* deletion leads to compromised fertility, but clearly many *Katna1*^-/-^ oocytes produce viable embryos that lead to healthy pups. Our investigations indicate that ovulation is not affected, suggesting fertility is reduced due to events that originate in oocytes and compromise subsequent fertilisation and embryo development. Surprisingly, we found MI spindles appeared normal, but abnormalities were detected in a proportion of MII spindles. In mammals, there is a remarkable reorganisation of microtubules required as the MII spindle forms immediately after Telophase I, and the data presented here suggests KATNA1 assists in this process. A small number of proteins have been identified that cause MII-specific effects including MAP kinase substrates MISS and DOCR1 (Lefebvre et al., 2002; Terret et al., 2003), as well as Cyclin A2 (Zhang et al., 2017). Further work is needed to understand if any of these proteins are involved in regulating KATNA1 function specifically in MII, where increasing evidence suggests specific mechanisms are necessary for MII spindle formation and function.

The disruption of spindle poles in a proportion of MII oocytes suggests, in the absence of centrioles, KATNA1 plays a role in coordinating the capture and organisation of microtubule organizing centers (ncMTOCs), that effectively act as the spindle poles in the barrel-shaped oocyte spindles. The requirement to harness MTOCs to form spindle poles in the short 90-minute MI to MII transition, relative to the 6-7-hour long prometaphase leading to MI spindle formation, may explain the specific susceptibility of MII to the loss of KATNA1. A role for katanin in mammalian oocyte meiosis is consistent with a conserved role in *C. elegans* where deletion of the *Katna1* orthologue, MEI1, also leads to disrupted meiotic spindles (Srayko et al., 2000; Srayko et al., 2006).

The increase in spindle abnormalities in MII stage oocytes appear to be the first phenotypic explanation for compromised fertility of conditional *Katna1*^-/-^ mice. The decrease in oocytes cleaving to the 2-cell stage after fertilization may be a result of chromosome segregation defects at exit from MII and/or in the first mitotic division, or alternatively, microtubules may play a role in the early events of fertilisation such as sperm head incorporation and pronuclear formation (Maro et al., 1986). The disrupted spindle organisation seen in KATNA1-deficient MII oocytes suggests an increase in aneuploidy underlies the decrease in fertility of conditional *Katna1*^*-/-*^ mice. Small litter size, as observed here, is consistent with aneuploidy being a key player in reduced embryo viability. Increased aneuploidy manifests in a phenotype over a broad time frame ranging from preimplantation arrest through to embryonic failure at implantation or beyond, depending on the specific chromosomes involved and the stage at which it aneuploidy happens (Hernandez and Fisher, 1999; Potapova et al., 2013). Furthermore, aneuploid blastomeres arising in cleavage divisions can be deleted by apoptosis (Singla et al., 2020) and aneuploid cells can be accommodated in the trophectoderm and resultant extra-embryonic tissues (James et al., 1995; Kalousek and Dill, 1983). The complexity of the potential origin and outcome of aneuploidy is also consistent with their being no single point of failure in KATNA1-deficient oocytes, and that the reduced fertility occurs across the developmental continuum from fertilization through to peri- and post-implantation development.

The maternal effect of *Katna1* deletion in oocytes was shown to be muted by the paternal allele introduced at fertilisation. Our experiments using parthenogenetic activation to create KATNA1-depleted preimplantation embryos, reveal that KATNA1 contributes to the mitotic cell divisions of preimplantation development. The 50% reduction in blastocyst formation, reduced cell number and increase in nuclear size point to failure of microtubule function in mitosis and cytokinesis. These effects are most likely attributed to abnormal spindle pole organisation, disrupted microtubule dynamics leading to spindle disruption and chromosome segregation errors, as well as disrupted midbody formation, some of which have been reported in the absence of KATNA1 in other systems (Buster et al., 2002; Gao et al., 2019; Jiang et al., 2017; Matsuo et al., 2013; Srayko et al., 2000; Srayko et al., 2006; Ververis et al., 2016).

The increased sensitivity of the embryonic mitotic divisions to KATNA1 depletion compared to the meiotic divisions may be associated with the transition from an acentriolar spindle in oocytes to a more typical mitotic spindle with centrioles and focused spindle poles by the 8-32 cell stage (Courtois et al., 2012; Gueth-Hallonet et al., 1993; Palacios et al., 1993). This is also consistent with findings in spermatogenesis where centriolar spindles persist and, unlike oocytes, katanin deletion has a major impact on spindle organisation and function (Dunleavy et al., 2022; Dunleavy et al., 2021; O’Donnell et al., 2012). Thus, the increasingly important role of katanin in preimplantation embryos relative to the oocyte may be a manifestation of the transition from an acentriolar to centriolar spindle organisation.

The main effect detected after depleting KATNAL1 during oocyte growth was a small but significant change in spindle size, an effect consistent with the role of katanins in spindle size scaling in Xenopus (Loughlin et al., 2011) and regulating spindle flux in Drosophila mitosis (Zhang et al., 2007). The differences were only 1-2μm, which is within 10% of the average spindle length in control oocytes. In the absence of other effects of spindle function these small differences are not expected to have any physiological significance and may be bought about by subtle changes in microtubule dynamics, as seen after depleting katanins from cell lines (Jiang et al., 2017; Zhang et al., 2011).

These subtle effects in our *Katnal1* genetic model are in stark contrast to the disrupted and multipolar spindles and meiotic arrest seen after acute knockdown of *Katnal1* using siRNA (Gao et al., 2019). This difference in severity of phenotype may be that the prolonged 36-44h cumulus-free culture period after siRNA transfection led to additional non-specific effects, or that it compromised oocyte quality, effectively sensitizing them to the effect of reduced KATNAL1. Interestingly, this appears consistent with our findings showing that in oocytes that failed to mature, it was only in the abnormal *Katnal1* knockout oocytes that we observed disrupted microtubule organisation in the form of excess cytoplasmic asters. Thus, in the absence of KATNAL1, poor quality oocytes may be more prone to amplifying abnormalities in microtubule function. Alternatively, there is increased evidence in a number of species that acute depletion of gene products leads to a more enhanced phenotype than genetic deletion *in vivo*, where compensatory mechanisms may come into play (El-Brolosy and Stainier, 2017; Tautz, 1992). In support of this, functional compensation of the katanin A-subunits has been documented (Dunleavy et al., 2022)

The presence of multiple catalytic katanin A-subunits in mouse oocytes may provide some level of functional redundancy so that the loss of any one of the family does not completely abolish fertility. This concept has been elegantly demonstrated in male germ cell development where the deletion of both *Katna1* and *Katnal1* subunits causes complete infertility, while *Katna1* deletion is without effect and *Katnal1* deletion leads only to subfertility (Dunleavy et al., 2022). Further, in *C. elegans* oocytes, where there is only a single Katanin A-subunit, and therefore no opportunity for functional compensation, deletion of the single subunit leads to failure of the meiotic cell divisions (Clark-Maguire and Mains, 1994; Loughlin et al., 2011; McNally and Thomas, 1998; O’Donnell et al., 2012; Srayko et al., 2000).

In summary, our study provides the first genetic evidence for a role of katanin in the production of viable fertile mammalian oocytes. Further studies examining the role of the regulatory subunit KATNB1, which typically demonstrates a more severe phenotype due to its regulation of multiple katanin A-subunits, will be interesting in the context of the oocyte to embryo transition in mammals.

## Experimental Procedures

### Mouse model generation and ethics

All animal procedures were approved by the Monash University School of Biological Sciences Animal Experimentation Ethics Committee and conducted in accordance with National Health and Medical Research Council (NHMRC) Guidelines on Ethics in Animal Experimentation.

Homozygous *Katna1*^*fl/fl*^ and *Katnal1*^*fl/fl*^ mice (C57BL/6J) were obtained from the laboratory of M. O’Bryan and crossed with *Zp3-Cre* transgenic mice (C57BL/6J; provided by E. McLaughlin, University of Newcastle, New South Wales, Australia) that specifically express *Cre* in growing oocytes to generate *Katna1*^*-/-*^ or *Katnal1*^*-/-*^ oocytes. To control for background strain effects on phenotype, *Katna1*^*Flox/Flox*^ and *Katnal1*^*Flox/Flox*^ mice were used as controls for *Katna1*^*-/-*^ and *Katnal1*^*-/-*^ mice respectively. For each strain, mouse genotypes were determined from tail biopsies using realtime PCR with specific probes designed for each allele (Transnetyx).

### Oocyte collection and culture

4–6-week-old C57BL6/J female mice were sacrificed for oocyte collection for all experiments described. For in vitro maturation, GV oocytes were collected in M2 (Sigma) containing 200 μM 3-isobutyly-1-methylx-anthine (IBMX). Culture was performed in drops of M16 medium (Sigma) under mineral oil (Sigma) at 37°C in a humidified atmosphere of 5% CO_2_ in air. For collection of MII stage oocytes, mice were superovulated by sequential intraperitoneal injections of 10IU pregnant mare’s serum gonadotropin (PMSG, Intervet) and 10IU human chorionic gonadotropin (hCG, Intervet) at timed intervals before collection.

### Fertilisation, parthenogenetic activation and embryo culture

Cauda-epididymis from 2 month old C57BL/6JxCBA F1 male mice were collected and sperm was allowed to capacitate in HTF media for 1 hour at 37°C in a humidified atmosphere of 5% CO_2_ in air. Cumulus mass collected from oviducts were then exposed to capacitated sperm and left to incubate for 2-4 hours before washing out excess sperm and transferring them to drops containing culture media. Pronuclei formation was checked 9h after fertilisation, 2-cell and blastocyst numbers were tabulated at 24h and 84h respectively. Superovulated MII eggs were briefly washed in hyaluronidase before placed into 10mM strontium chloride calcium-free CZB media supplemented with 1μg/ml cytochalasin B.

### Immunofluorescence and live-cell imaging

Oocytes or embryos were fixed in a solution of 4% paraformaldehyde and 2% triton X-100 in PBS for 30 min at room temperature. Blocking was performed in PBS with 10% goat or donkey serum, 10% BSA and 2% Tween for 60 min at room temperature. Antibodies used for immunofluorescence-labelling: FITC-conjugated mouse anti-α-tubulin (1:200, Alexa Fluor 488, Invitrogen). DNA was labelled using 10 min incubation in Hoechst 33342 (10 μg/ml, Sigma-Aldrich). Serial Z sections of fixed oocytes/eggs were acquired at room temperature in a glass-bottomed dish using laser-scanning confocal microscope imaging system (SP8, Leica). For embryo live-cell imaging, 2-cell embryos were incubated and imaged for 4 days using the ESCO Medical Miri TL. The microinjected oocytes were incubated in M2 media at 37°C under mineral oil and imaged with Leica SP8 confocal microscope.

### Reverse transcription PCR

RNA was extracted and purified from GV oocytes using the QIAgen RNAeasy Plus Micro Kit. Reverse transcription was carried out using the High Capacity cDNA reverse transcription kit (Applied Biosystems). Polymerase chain reaction was then performed on the cDNA using the Bioline MyTaq Red Mix using the following primers for Katna1 (5’-ACT GTG TGG TCA TCT GCT G-3’ 5’-TAA ATA TGG TAG CTG GGG AGT-3’) Katnal1 (5’-CCT GAC TTC CCC GTG TCT TG-3’, 5’-GTG ACA AAC CTG CAA GTC GG-3’), and β-actin (5’-TGA GAG GGA AAT CGT GCG TGA CA-3’ 5’-CCA GAG CAG TAA TCT CCT TC-3’). PCR conditions is as follows, 1 cycle 95°C for 60s, 34 cycles of 95°C for 15s, 60°C for 15s, 72°C for 20s, 1 cycle of 72°C for 120s. PCR product was then run on a 2% agarose gel at 120V for 1h, then imaged on the Biorad Geldoc.

### Data and statistical analysis

Z-stack immunofluorescent images of blastocysts were imported into Imaris for processing to determine nuclei volume. Border of each nucleus was drawn out individually on every Z-stack layer to create objects that could be quantified using the computer’s algorithms.

Statistics was performed using student’s t-test, or Chi^2^ test with Sigma Plot or GraphPad Prism Software. Analysis was performed for at least three replicate experiments and represented as means and SEMs, unless otherwise stated. Differences at p< 0.05 were considered significant. Level of significance is denoted by n.s. p>0.05, * p<0.05, **p<0.01, ***p<0.001, ****p<0.0001.

## Acknowledgements

We thank Dr Greg Fitzharris for his contributions to the project. We thank Monash Microimaging and Monash Animal Research platform for the use of their facilities and services.

---

**Supplementary Figure 1.**
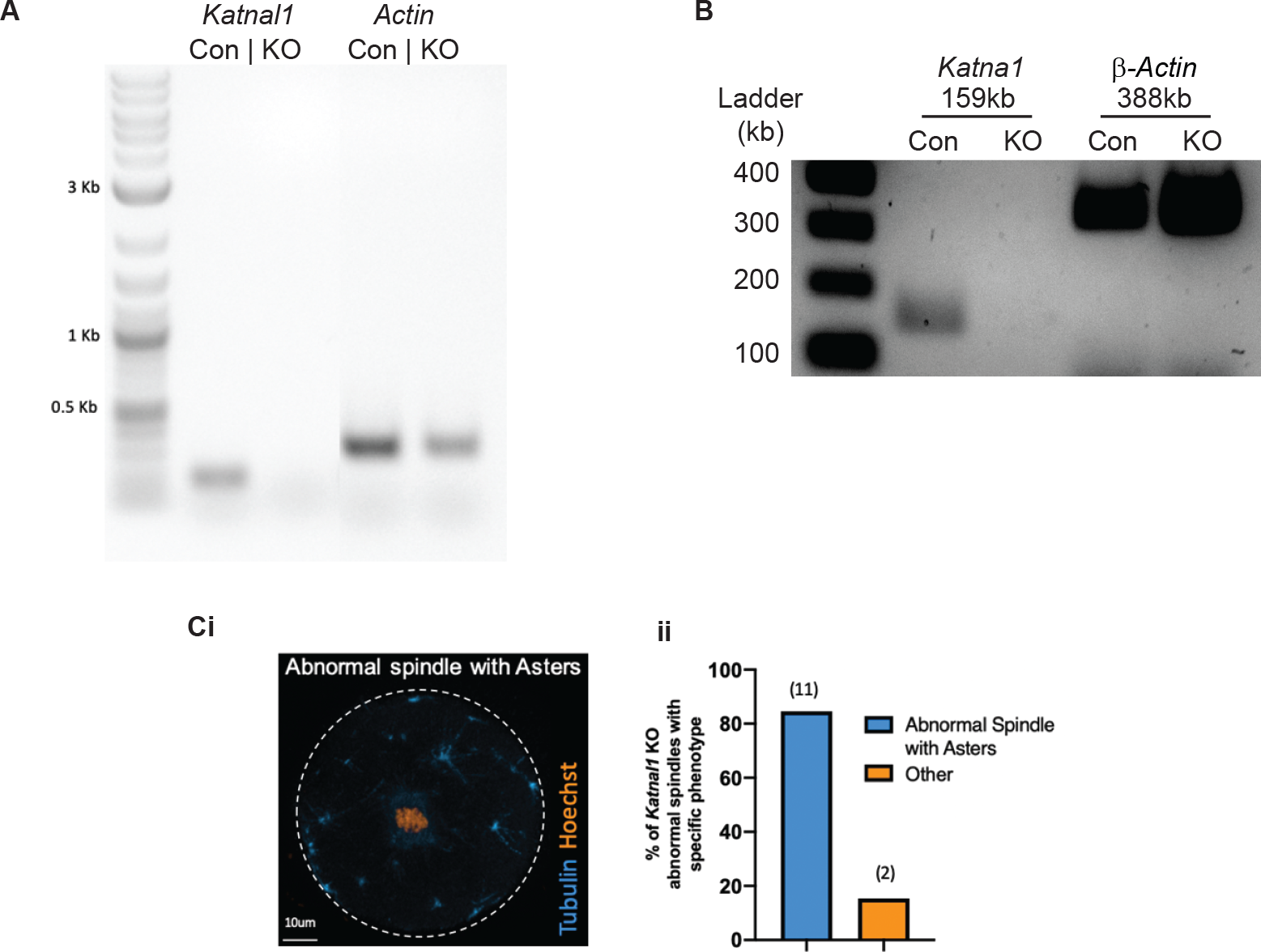
Katna1 and Katnal1 knockout RT-PCR (A) Knockout of Katnal1 mRNA shown with RT-PCR using β-actin as an experimental control. (B) Knockout of Katna1 mRNA shown with RT-PCR using β-actin as an experimental control. (Ci) Representative image of abnormal Katnal1-/- MI oocytes displaying asters throughout cytoplasm. (Cii) Percentage of abnormal Katnal1-/- MI oocytes displaying asters throughout cytoplasm.

